# A Multi-Scale Computational Model of the effects of TMS on Motor Cortex

**DOI:** 10.1101/064337

**Authors:** Hyeon Seo, Natalie Schaworonkow, Sung Chan Jun, Jochen Triesch

## Abstract

The detailed biophysical mechanisms through which transcranial magnetic stimulation (TMS) activates cortical circuits are still not fully understood. Here we present a multi-scale computational model to describe and explain the activation of different cell types in motor cortex due to transcranial magnetic stimulation. Our model determines precise electric fields based on an individual head model derived from magnetic resonance imaging and calculates how these electric fields activate morphologically detailed models of different neuron types. We predict detailed neural activation patterns for different coil orientations consistent with experimental findings. Beyond this, our model allows us to predict activation thresholds for individual neurons and precise initiation sites of individual action potentials on the neurons’ complex morphologies. Specifically, our model predicts that cortical layer 3 pyramidal neurons are generally easier to stimulate than layer 5 pyramidal neurons, thereby explaining the lower stimulation thresholds observed for I-waves compared to D-waves. It also predicts differences in the regions of activated cortical layer 5 and layer 3 pyramidal cells depending on coil orientation. Finally, it predicts that under standard stimulation conditions, action potentials are mostly generated at the axon initial segment of corctial pyramidal cells, with a much less important activation site being the part of a layer 5 pyramidal cell axon where it crosses the boundary between grey matter and white matter. In conclusion, our computational model offers a detailed account of the mechanisms through which TMS activates different cortical cell types, paving the way for more targeted application of TMS based on individual brain morphology in clinical and basic research settings.

## Introduction

Transcranial magnetic stimulation (TMS) is a neurostimulation and neuromodulation technique that noninvasively activates neurons in the brain^1,2^. It generates a time varying magnetic field using a coil above the head, which induces an electric field in the brain that can be of sufficient magnitude to depolarize neurons. In recent years, TMS has been widely tested as a tool for diagnosis and treatment for a broad range of neurological and psychiatric disorders^3–5^. Although the efficacy of TMS has been demonstrated, there remains a large degree of uncertainty regarding the factors influencing the affected brain areas and relevant circuits.

To provide a better understanding of the biophysical mechanisms behind TMS, several computational studies have been performed to try to reveal the effects of a number of parameters that lead to variable outcomes. The majority of models predicts the brain regions influenced by TMS based on stimulus-induced electric fields^6,7^. While early studies utilized spherical models of the human head, in recent years high-resolution volume conduction models of the head were developed from human magnetic resonance imaging (MRI) to the improve accuracy of calculated electric fields^8–15^. These models revealed that the geometry of the volume conduction model, such as complex gyral folding patterns, is one of the key parameters determining the induced electric field. In addition, computational studies were extended by connecting numerical results with experimental observations to show the correlation between computed electric fields with physiological observations^16–18^.

Directly monitoring target cells’ activities under stimulation would be immensely valuable for the interpretation of TMS effects, but few such studies exist^19,20^. However, computational studies can explore the effects of the electromagnetic fields on neural activation by simulating models of neural stimulation *in silico*. In early computational models, straight axonal fibers were considered numerically and the response of neurons induced by the external field was modeled by means of the cable equation^21,22^. Later models investigated the role of cell morphology using multi-compartmental modeling^23–25^. Since the responses of cortical neurons vary depending on not only the neuronal morphology but also orientation relative to the induced electric field and stimulus amplitude^6,26,27^, anatomical information such as cortical folding that induces a wide range of field orientations was fed into the neuronal models by applying the calculated electric field from the head model to the neuronal models^28,29^.

Here, we use an advanced multi-scale modeling approach that combines a high-resolution head model with detailed multi-compartmental neuron models. We construct an anatomically realistic head model based on MRI and calculate the external currents that affect neurons via the TMS-induced electric field with high accuracy. We concentrate on the hand knob area of the motor cortex that is the predominant target of many TMS studies^16,30^. A multitude of layer 5 and layer 3 pyramidal neurons (L5/L3 PNs) is incorporated on the basis that they might be primary activators of the corticospinal tract and provide the main input to the direct pathway^24,28,31,32^. We estimate the target area of activation as a function of coil orientation as well as the stimulation intensities required to activate neurons. Finally, we predict the precise sites where the neurons initiate their action potentials.

## Methods

In order to study the cellular effects of TMS in the brain we employed a multi-scale computational modeling approach combining a volume conductor head model with detailed neuronal models of cortical pyramidal neurons. The motor cortex, especially the hand area, was considered as a cortical target location. The volume conductor head model was used to simulate the stimulus-induced electric fields. The precise impact of these fields on different neural targets was evaluated using multi-compartmental models of pyramidal neurons embedded into the head model. This allowed us to predict differences in individual neuron’s susceptibility to TMS depending on neuron placement and coil orientation.

### Volume conductor model

The simulated effects of TMS depend not only on stimulation parameters but also on the anatomical information specified in the volume conductor model. To calculate the precise electric field, a volume conductor head model for TMS that reflected T1-weighted and T2-weighted magnetic resonance (MR) images was constructed using SimNibs^14,33^. Briefly, segmentation of white matter (WM), gray matter (GM), cerebrospinal fluid (CSF), skull and skin was based on FreeSurfer^34,35^, FSL^36^ and MeshFix^37^, as shown in Fig 1(a). Then, the head model was constructed by generating an optimized tetrahedral volume mesh using an enhanced resolution in the region of interest (ROI) around the hand knob using Gmsh^38^. The total number of tetrahedral elements was approximately 5.6 million. At each layer of the head model, isotropic conductivity was assigned with the following values (in S/m): WM: 0.126; GM: 0.276; CSF: 1.654; skull: 0.01; and skin: 0.465.

**Figure 1.**
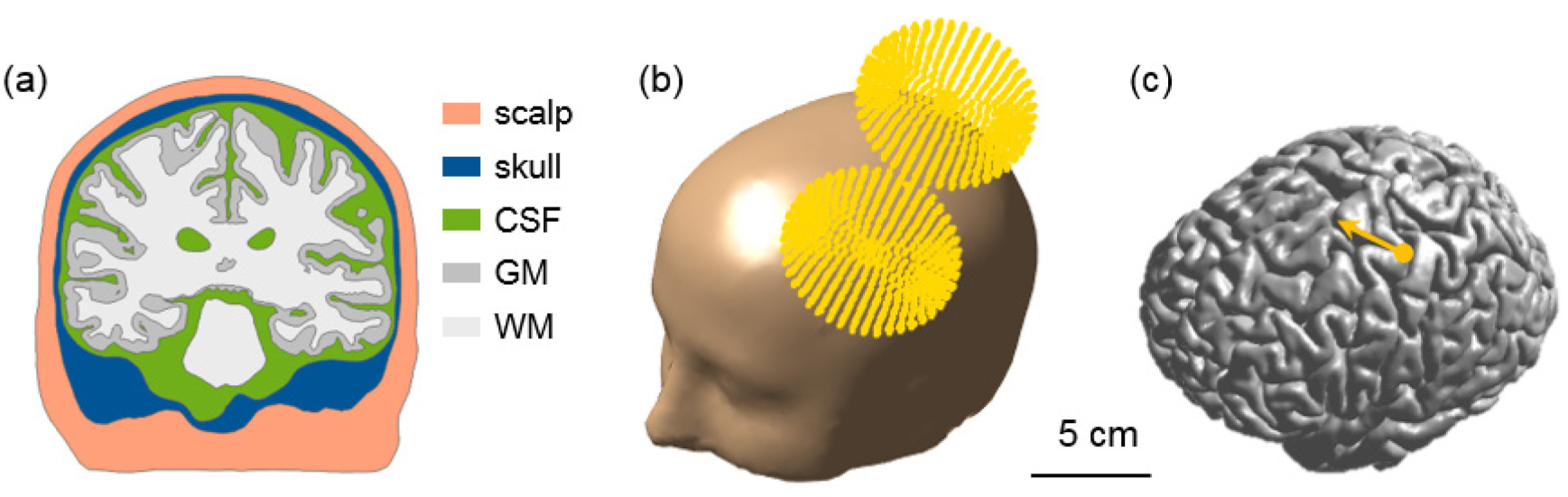
The volume conductor model and coil placement. (a) Cross-section displaying the scalp, skull, cerebrospinal fluid, gray matter and white matter. (b) The computed coil location is superimposed on the head model. (c) The yellow dot indicates the location of the center of the TMS coil on the border between gray matter and cerebrospinal fluid, and the coil handle is oriented in the direction of the yellow arrow.

### Field calculations

The electric field induced by TMS, 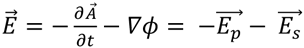, consisted of primary 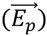 and secondary 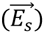 electric fields. The primary electric field was directly determined by the coil geometry and the head model. The secondary electric field was solved via a finite element method using the GetFEM++ library and MATLAB^33,39^. The Magstim 70 mm figure-8 coil was represented by magnetic dipoles positioned above the hand knob (Fig. 1) and the stimulator output was set to 1 A/µs. The coil orientation was defined relative to the direction of the central sulcus such that the electric field induced was in anterior to posterior direction (Fig. 1(c)). Then, three additional coil orientations were tested by rotating in steps of 45 degrees and reversed orientations were simulated by changing the sign of the current through the coil.

To investigate the TMS-induced cellular effects, we quantified the magnitude of the electric field 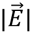 and the orthogonal component of the electric field 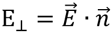 to the gray matter surface, where 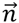 is the normal vector for the boundary surface element. The orthogonal component was expected to contribute to TMS-induced brain activation by the theoretical *cortical column cosine model of TMS efficacy* (*C*^3^-*model*)^10,40^.

### Multi-compartmental neuronal models

We adapted existing multicompartmental models of layer 5 and 3 pyramidal neurons (L5/L3 PNs) from cat visual cortex^41^ using the NEURON simulation software^42^. The electrical properties were unchanged from the original models. Briefly, a high density of fast, inactivating voltage-dependent Na^+^ channels were present in the axon hillock and axon initial segment, and a low density of these channels was presented in the soma and dendrites. Slow Ca^2+^-dependent K^+^ channels and high threshold Ca^2+^ channels were located in the soma and dendrites. Except for the dendrites, fast K^+^ channels were present. L5/L3 PNs were combined virtually with the head model and modified to accommodate the irregular geometry of the cortex^28,29,43–46^, as shown in Fig. 2. The dendritic trees were lengthened or shortened by re-scaling them according to the local dimensions of the cortex such that dendrites reached layer 1 and the orientation was perpendicular to the cortical surface^45,47^. Since the morphology of the dendritic trees was not symmetric and it might influence the neuronal activation, L5/L3 PNs had randomly rotated dendritic trees at different locations. The axons of L5 PNs were defined to curve beyond the boundary between GM and WM toward the corpus callosum. The axons of L3 PNs were defined to terminate in layer 5/6 within the GM. To reduce superfluous computations, we preselected a region of interest (ROI) of 50 × 50 × 50 mm^3^ around the hand knob and then placed L5/L3 PNs in each triangular element comprising the gray matter surface. Altogether, a total of 10,888 L5 PNs and 10,888 L3 PNs was constructed. This process was implemented in MATLAB (MathWorks, Natick, MA, USA).

**Figure 2.**
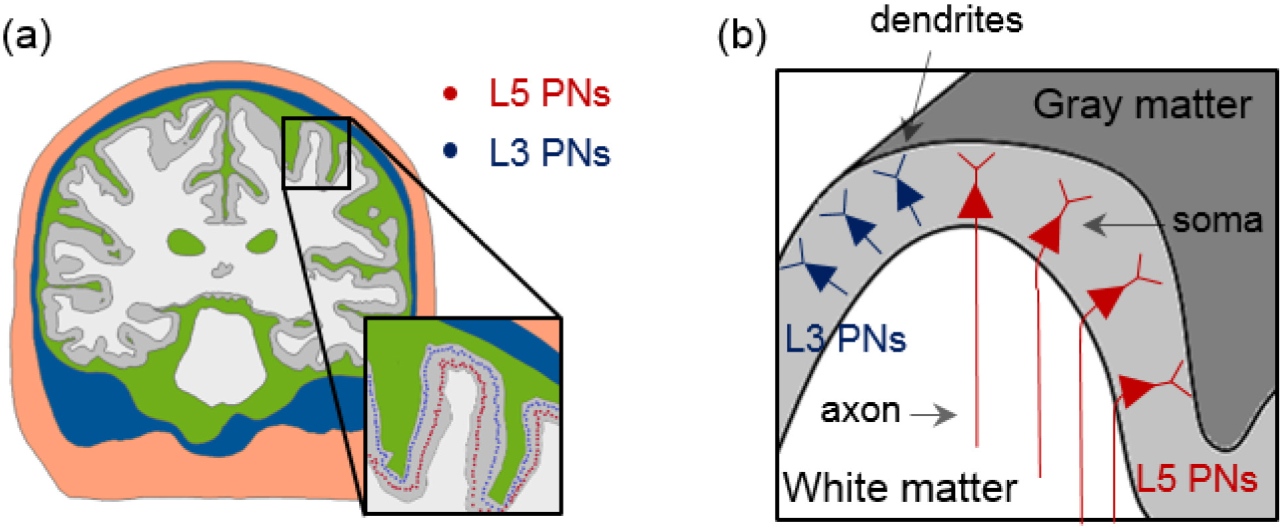
The placement of L5/L3 PNs in the head model. (a) The distributions of somata of L5/L3 PNs are marked as colored dots (red: L5; blue: L3). (b) A schematic view of the distribution of the L5/L3 PNs is shown along the cortex folding (gray colored area); note the bending of L5 PN axons when crossing the boundary between gray matter and white matter.

### Computation of neuronal activation induced by stimulation

The membrane potentials induced by stimulation were approximated by adding an external current source *I_ext_* to the cable model^2,21,22,24,25^:

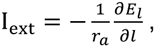

where r_a_ is the axial resistance per unit length and E_l_ represents the component of the electric field that is parallel to each compartment of the PNs. The derivative of the electric field along each compartment was calculated at each center point by 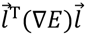, where ∇*E* contains the components of the electric field gradient tensor that are estimated by computing the difference of electric fields at neighboring points displaced by ±1 mm along each axis^48^.

We calculated a monophasic pulse that induced a fluctuating magnetic field through an RLC circuit as detailed in^49^,

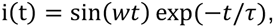

where w = 30 kHz is the angular frequency and τ = 0.08 ms is the decay time. The *I_ext_* at each compartment was then multiplied by the normalized time derivative of the monophasic pulse. Finally, we obtained the spatial and temporal membrane potential dynamics. They were used to measure the excitation thresholds, the stimulation site and action potential propagation.

## Results

Fig. 3 depicts the magnitude of the electric fields (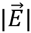, top row) and the orthogonal component of the electric fields (E_⊥_, bottom row) for different coil orientations. All calculations were performed for a rate of change of the coil current of 1 A/µs. Electric fields had higher magnitudes in the precentral and postcentral gyrus and focused on the top of the gyri, regardless of coil orientations. We observed only slight changes in the field strengths depending on coil orientation. In contrast, the orthogonal component of electric fields (E_⊥_) showed different spatial patterns compared to the electric field magnitude. High strengths of E_⊥_ were found on the walls of the gyri and strongly depended on coil orientation. Furthermore, while the spatial extent of 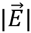 was the same for the standard orientation and the +180 E_⊥_ in the degree orientation, the sign of +180 degree orientation was reversed due to the reversed sign of the induced electric fields. Interestingly, the maximum value of 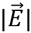 depended on coil orientation; its was lowest in the standard coil orientation and highest at +135 degrees. However, the maximum values of E_⊥_ were highest for the standard coil orientation and lowest at +90 degrees.

**Figure 3.**
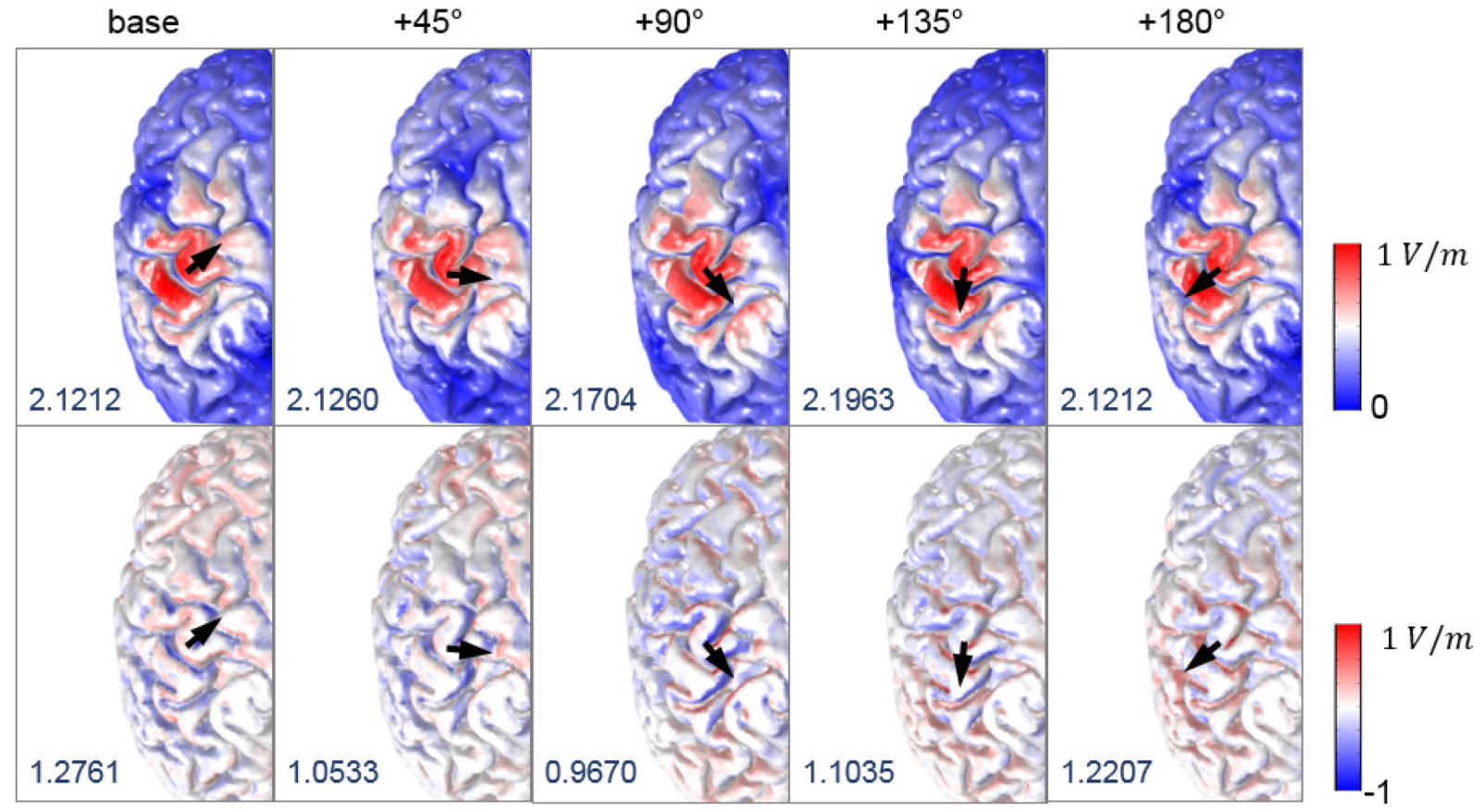
Effects of coil orientation on the electric fields. The spatial patterns of magnitude of electric fields (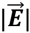, top row) and its component orthogonal to the gray matter surface (**E**_⊥_, bottom row) are visualized; the color scale is adapted for better visualization. The black arrows indicate different coil orientations, and the maximum value of 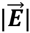 and **E**_⊥_ (measured in V/m) are given in the bottom left of each figure.

To assess the neuronal activations as a function of coil orientation, we determined the excitation threshold required to cause action potentials of L5/L3 PNs. For each coil orientation, we kept increasing the stimulator output until a neuron generated an action potential or we reached a maximum rate of change of the current defined as 171 A/µs. Our focus is on the excitability for a stimulation intensity corresponding to 67 A/µs, as this value corresponds to the average motor threshold for the Magstim 200 stimulator connected to the coil modeled^18,29,48,50^. The excitability of L5/L3 PNs was predicted either by the direct estimation of the electric field (E_⊥_map in the Fig. 4(c)) or by simulating the induced depolarization and firing of the detailed neuronal models (threshold maps in Fig 4(d,e)). The color of the threshold maps represents the stimulator output necessary to activate the corresponding cell and the estimated excitable area in the E_⊥_ maps. The blue colored areas indicate an excitability in the opposite direction, because the head model was linear with respect to the electric field. As shown in Fig. 4(a), we virtually divided the precentral and postcentral gyrus to better visualize the excitability in the walls of the gyrus.

**Figure 4.**
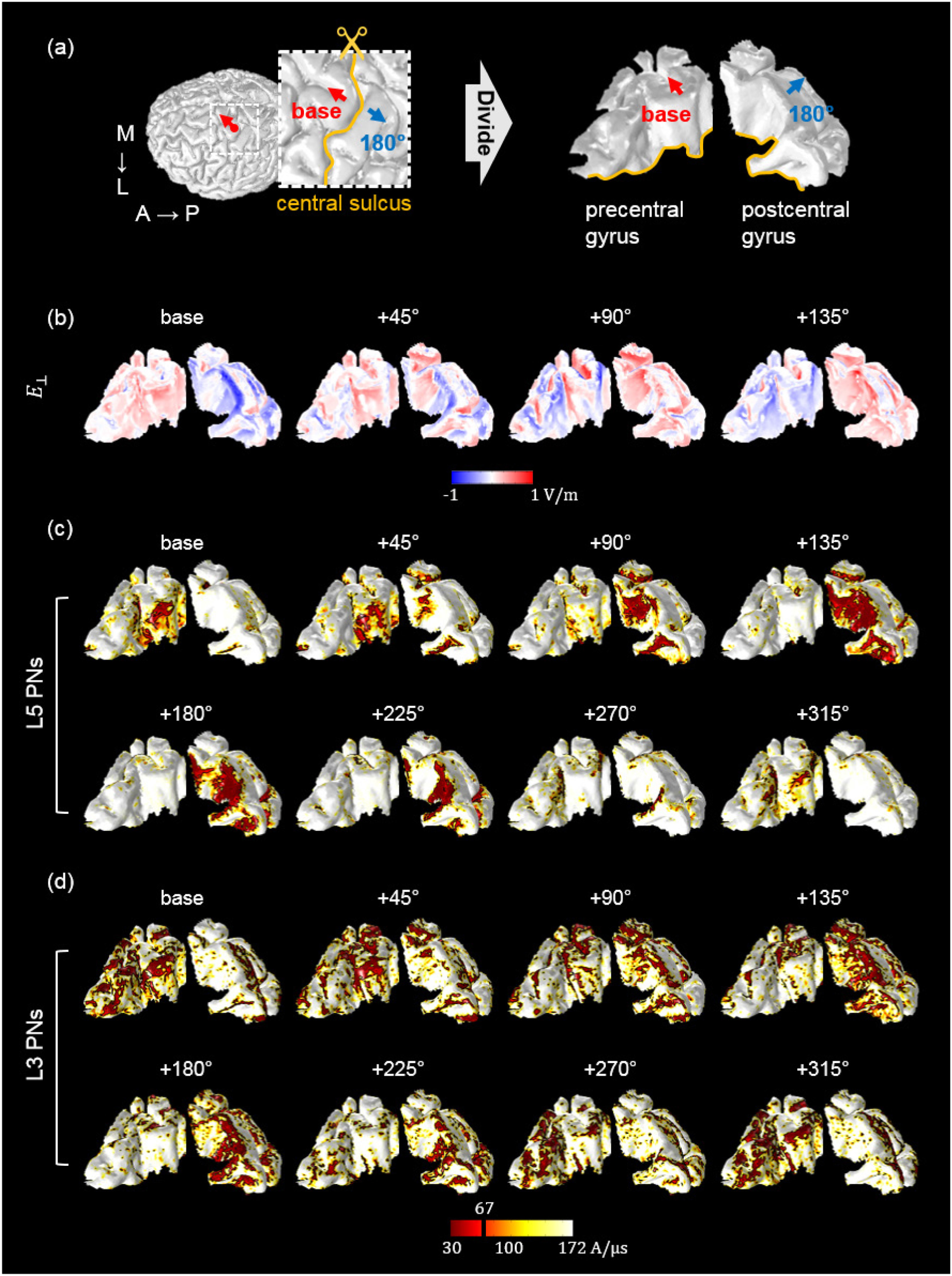
The spatial extent of predicted excitability based on the orthogonal component of the electric field (**E**_⊥_) and detailed simulations of L5/L3 PNs. (a) The red dot on the border between GM and CSF indicates the location of the center of the coil. The base orientation is shown as red arrows. The inset represents the region of interest in which PNs were distributed. The blue arrows indicate the opposite coil orientation (+180°). The precentral and postcentral gyri were virtually divided for visualization purposes. The spatial patterns of **E**_⊥_ (b) and threshold maps of L5 (c) and L3 (d) PNs depended on coil orientation as shown. The black and red colored areas in the threshold maps (c-d) indicate the excitable areas under the stimulator output corresponding to the average motor threshold (**67 A/μs**). The directions of coil orientations in the 2^nd^ row are the opposite directions of the 1^st^ row (in the threshold map in (c and d)) simulated by changing the sign of the current through the coil. Note how the excitable areas strongly depend on the coil orientation.

In L5 PNs, we observed that the predicted excitability depended on coil orientation for both E_⊥_ and threshold maps (Fig. 4(c,d)). For the base orientation and +45 degrees, a high excitability was predominantly observed in the wall of the precentral gyrus. In contrast, for orientations +90 degrees to +225 degrees we observed high excitability in the wall of the postcentral gyrus. Comparing these threshold maps to the E_⊥_ maps, we see that the threshold maps show activated L5 PNs in some additional smaller areas with comparatively small E_⊥_ values. From +90 to +180 degrees, the excited regions were quite well matched to the results from E_⊥_. Furthermore, in the standard direction, the spatial extents of L5 PNs that were activated for stimulation intensities corresponding to the motor threshold seemed to enlarge with increasing coil rotation, while for the opposite direction of the coil current highly excitable areas shrank with increasing coil rotations.

Overall, the excitability in L3 PNs showed behavior comparable to that of L5 PNs (Fig. 4 (d,e)), but notable differences in threshold maps between L5 and L3 PNs were as follows: while L5 PNs in the top of the gyri were never activated, L3 PNs were excited in the top and also the wall of gyri. The excitable areas of L3 PNs caused by the +90 and +135 degree stimulations were relatively focused on the upper parts of the wall of the postcentral gyrus, while L5 PNs placed in the deeper parts of the sulcus were activated. Furthermore in L3 PNs, the excitability in the precentral gyrus and the postcentral gyrus was comparable and a bigger area was affected than for L5 PNs. The discrepancies between L5 and L3 PNs confirmed that the morphology and placement of neuronal models has an important impact beyond the position relative to the coil.

The percentage of excited neurons for a stimulation intensity at the motor threshold is shown in Fig. 5. Consistent with the results for the threshold maps in Fig. 4 we found that the percentage of excited L5 PNs increased from base to +135 degrees and then decreased gradually thereafter. Similarly, the maximum percentage of excited L3 PNs was observed when the coil was rotated at +135 degrees and L3 PNs had about 2 times higher activations over all coil orientations. The overall percentage of excited neurons under the maximum stimulator output (171 A/µs) had a similar pattern; in standard orientation, 29% and 38% of L5 and L3 PNs were activated, respectively, and these numbers increased to 33% and 44% when the coil was oriented at +135 degrees.

**Figure 5.**
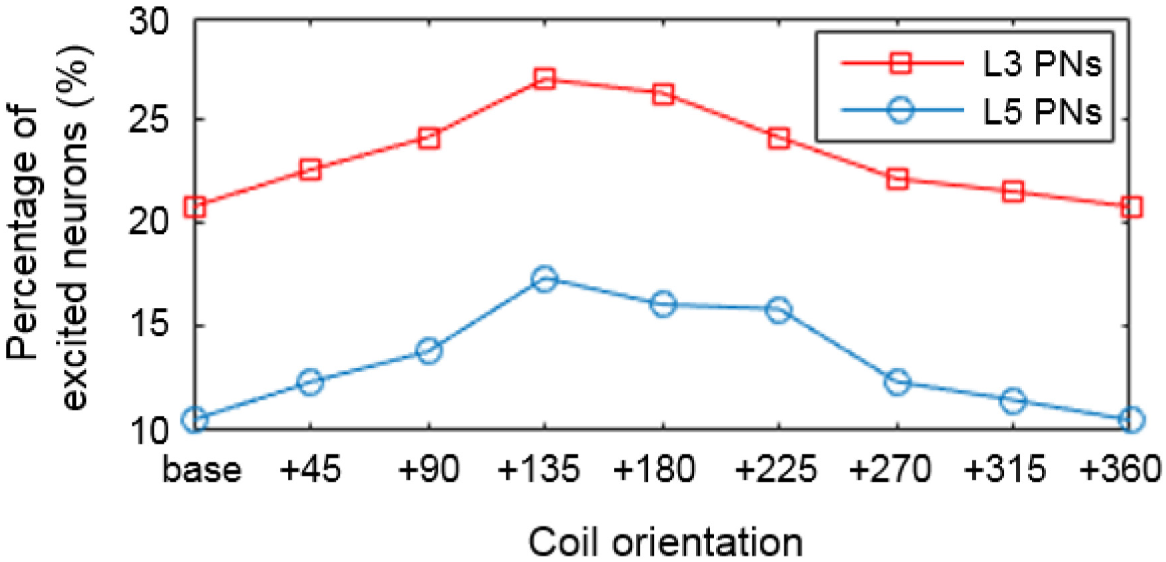
The percentage of L5 and L3 PNs that are activated for a stimulation intensity at the motor threshold (**67 A/μs**) as a function of coil orientation.

The majority of action potentials were initiated at the axon initial segment and others at the axon near the boundary between GM and WM for L5 PNs and at the middle and terminal points for L3 PNs (Fig. 6). In the base orientation, threshold stimulation elicited action potentials first at the initial segment for 90% of both the L5 and the L3 PNs. This fraction increased with increasing coil rotations up to 97% at +135 degrees in L5 PNs and up to 95% at +90 degrees in L3 PNs. Example plots of membrane potential dynamics induced by the threshold stimulus evoking action potentials are shown in Fig. 7. We observe the propagation of the action potentials from the axon initial segment to the more distal parts of the neurons. In both L5 and L3 PNs, following the action potential at the initial segment, the soma was activated as it is closest to the initial segment. The terminal points of the axons were activated last as they are most distal from the axon initial segment. Since the axon of a L5 PN is quite long compared to that of a L3 PN, the arrival of the action potential at the terminal point was substantially delayed. Similarly, dendrites of L5 PNs showed delayed activation while in the L3 PNs dendrites were occasionally activated early.

**Figure 6.**
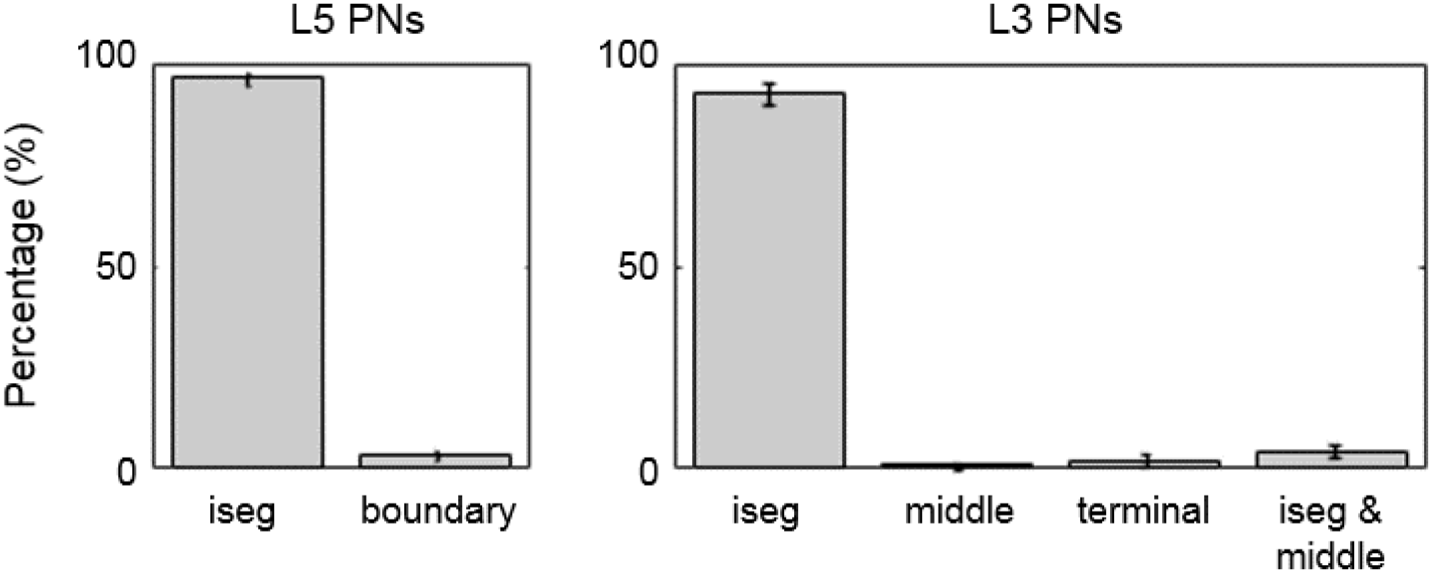
The percentage of action potential initiation sites of L5 and L3 PNs for a stimulation intensity corresponding to the motor cortex threshold (**67 A/μs**) averaged over different coil orientations. Sites include the axon initial segment (iseg) and the boundary between gray matter and white matter (boundary). Additionally, the terminal part (terminal) and middle point of the axon (middle) for L3 PNs were considered. Most action potentials are first evoked at the axon initial segment of L5 PNs (96.31±1.72%) and L3 PNs (92.76 ±2.42%). The remaining number of L5 PNs show action potential initiation at the axon near the boundary between gray matter and white matter. Only few L5 PNs (0.49±0.14%) initiate action potentials simultaneously at the axon initial segment and the GM-WM. For L3 PNs, middle (1.05±0.78%) and terminal points (2.08 ±1.45 %) of axons are also activated occasionally.

**Figure 7.**
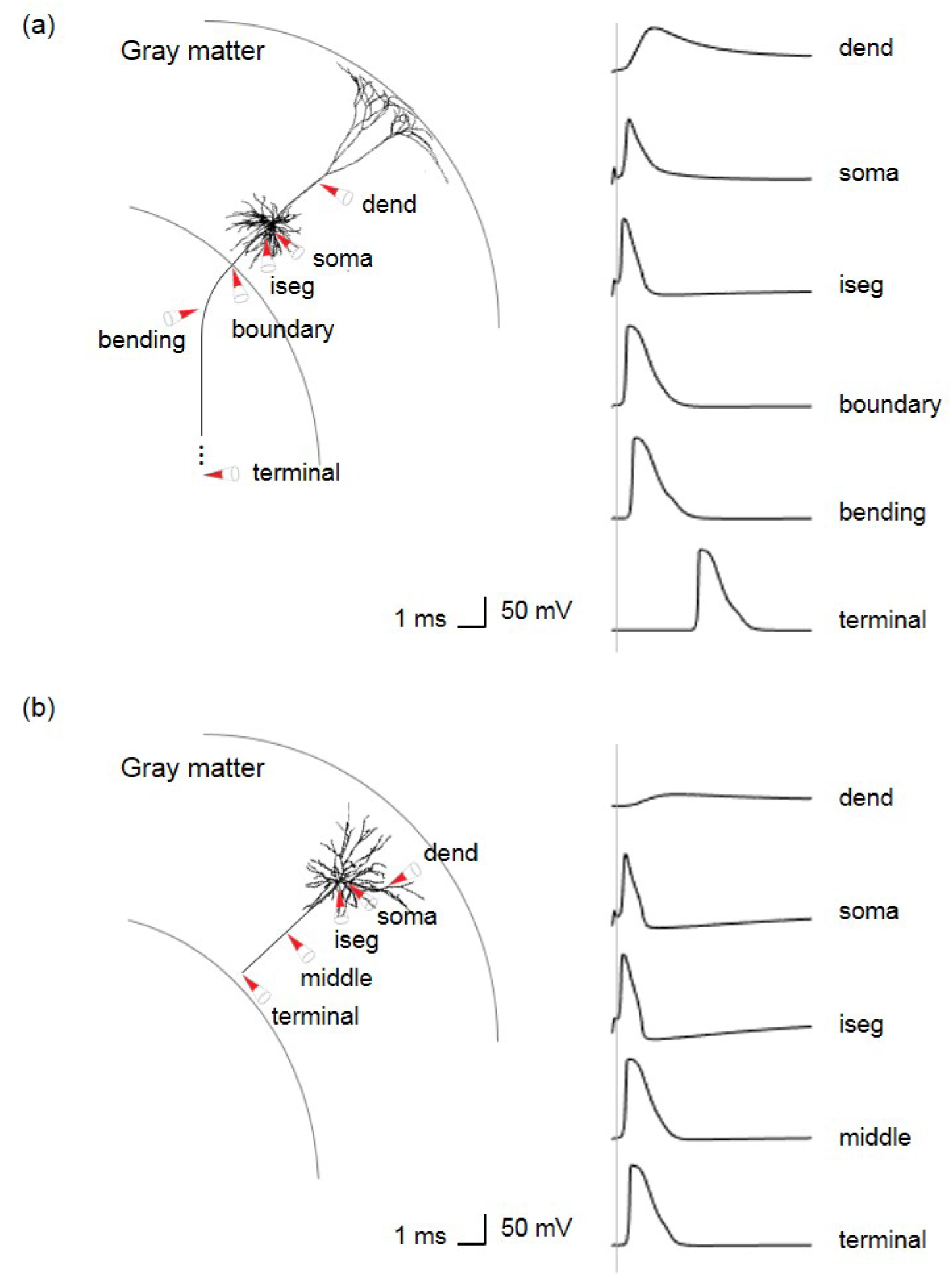
Virtual recordings of membrane potential dynamics of L5/L3 PNs. The simulated recordings were performed from dendrites (dend), soma and parts of the axons, as indicated by the red colored cones. (a) In a L5 PN, the membrane potentials are recorded at the axon initial segment (iseg), the location where the axon crosses the boundary between gray matter and white matter (boundary), bending and terminal points. (b) Additionally, the middle points of axons of L3 PNs are recorded.

The PNs that were morphologically reconstructed had asymmetric dendritic trees that might affect the neuronal responses. We studied the impact of dendritic trees on threshold maps and the percentage of excited neurons for a stimulation intensity at the motor threshold by rotating them in steps of 45 degrees around the axis defined by their apical dendrite for a fixed coil orientation at +180 degrees. In the threshold maps for L5 PNs as shown in Fig. 8, the highest variations of the thresholds caused by these rotations were observed in the boundary between the top of the postcentral gyrus and the sulcus. However, the coil dependency in the threshold maps did not change and thus the L5 PNs toward the postcentral gyrus were activated consistently. Compared to the percentage of excited L5 PNs with randomly rotated dendritic trees (16.12% as shown in Fig. 5), the fixed orientation of dendritic trees induced changes in the fraction of activated neurons of up to 2%. The threshold variations in the L3 PNs were hardly noticeable compared to those of the L5 PNs. The percentage of excited L3 PNs was 26.4% with randomly rotated dendritic trees, and when the orientations of dendritic trees were fixed it resulted in changes of at most 0.3%. Thus, we found that the morphology of the dendritic tree of the L5 PN model had a bigger impact than that of the L3 PN model, possibly due to its greater lack of rotational symmetry. Overall, however, rotations of the dendritic trees around the axis defined by the apical dendrite did not alter the spatial extent of activated regions much.

**Figure 8.**
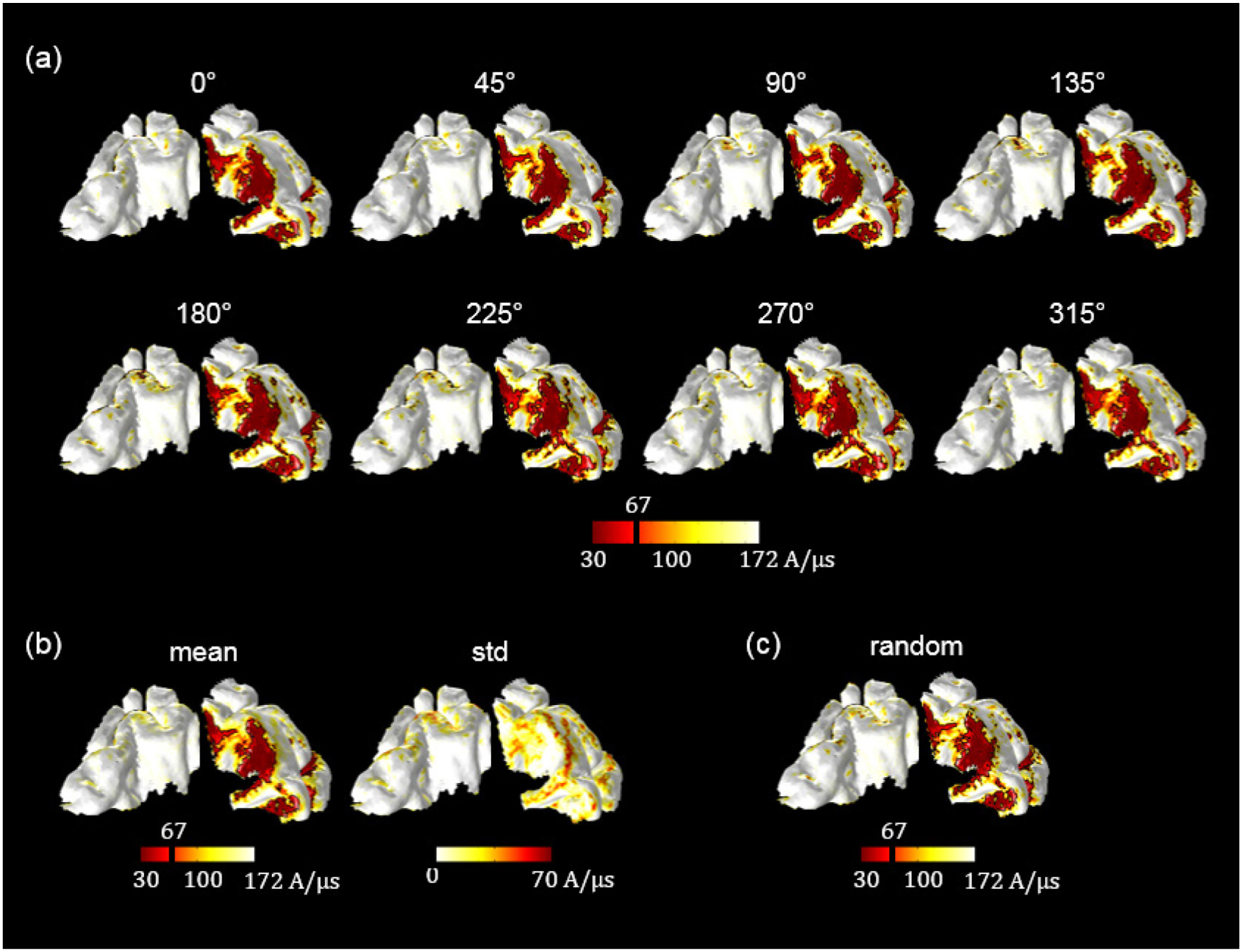
The impact of dendritic tree morphology of L5 PNs was evaluated by rotating them in steps of 45° for a fixed coil orientation at +180°. (a) The threshold maps according to the different orientations of dendritic trees and (b) its mean and standard deviation are shown. The map of standard deviations in (b) indicates precise orientation of the dendritic tree can alter activation thresholds in a noticeable fashion in certain situations. Overall, however, the activated areas in (a) do not change much compared to the threshold map for randomly rotated dendritic trees in (c).

## Discussion

The detailed mechanisms through which TMS activates cortical cells and cortical circuits are still not fully understood. In this study, we used multi-scale computational modeling to predict cortical activation as a function of coil orientation in two different ways. First, we simply considered the strength of the component of the TMS-induced electric field that is orthogonal to the gray matter surface as suggested by the C^3^-model^10,40^. Second we developed a detailed computational modeling approach that combined an anatomically realistic head model with complex multi-compartment neuronal models of L5/L3 PNs and quantified their stimulation thresholds. A major finding was the characterization of the induced electric fields and the thresholds of L5/L3 PNs as a function of coil orientations as shown in Fig 4. In addition, threshold variations according to different morphologies of PNs were observed.

The magnitude of the electric field was considered first, because the strength of the electric field is commonly used to extrapolate neuronal activation^6,7,51^. We found that the magnitude of the TMS-induced field is focused on the top of the gyrus, which is in agreement with previous modeling studies^10,12–14^. However, the electric field magnitude showed little dependency on coil orientation^10^. Then, we investigated the directional electric field, especially the orthogonal component that is perpendicular to the cortical surface, as this has been suggested to contribute most to the TMS-induced activation according to the C^3^-model^10,40,52^. We found a strong dependence of the orthogonal field component on coil orientation, as shown in Fig. 3. In contrast to the electric field magnitude, the highest field values were found in the sulcal walls and never on the apex (or crown) of the gyrus.

While the analysis of TMS-induced electric fields has been widely addressed in the past, the incorporation of multi-compartment neuronal models has hardly been investigated. To permit a more detailed understanding of the biophysical mechanisms of TMS, a few previous modeling studies employed detailed neuronal models and calculated the membrane potential dynamics generated by the electromagnetic field. However, these attempts had various limitations. First, in early studies no anatomical information on large-scale brain morphology was applied^23–25^. Rather than constructing a finite element head model, these studies applied a uniform electric field to the neuronal model. Even though such investigations achieved reasonable results regarding the neuronal activation, they did not consider the effects of the complex folding patterns of the cortex and the effects of tissue borders such as the borders between GM on the one hand and CSF or WM on the other hand. However, the importance of anatomically realistic head models has been shown convincingly^12–14^. Furthermore, the impact of detailed brain anatomy has been considered in various methods of brain stimulation and substantial differences have been demonstrated by improving anatomical information related to the head model^53–56^. Salvador et al. (2011) investigated neuronal responses using a simplified head model of a cortical sulcus with several types of neurons and found changes of the stimulation threshold depending on the pulse waveform and the coil orientation^29^. However, the used head model had an approximated geometry restricted to the motor cortex and a full complex geometry, such as the hook-shaped hand knob, was not considered. Furthermore, the modeled coil orientation was limited to anterior to posterior and its reversal due to the simplified geometry of the head model. Most recently, Goodwin and Butson (2015) proposed a more realistic approach that integrates an anatomically realistic head model derived from MR images with detailed neuronal models^28^. They considered the excitability of neurons as a function of coil orientation. However, in contrast to our results, excitability maps hardly showed a systematic dependence on coil orientation and activation thresholds were lower in the gyral crown. We speculate that this might be caused by the different morphology of PNs or the different way in which they calculated the external currents to simulate neuronal responses. Also, we considered two types of L3/L5 PNs spread over a wider region of the cortex. Finally, we also established the site of action potential initiation and found that most PNs are activated at the axon initial segment and action potential initiations at other parts of the neuron are comparatively rare.

The threshold maps we calculated demonstrate acute sensitivity to coil orientation, but different spatial extents were observed according to the different morphologies of the PNs. In L5 PNs, activation thresholds were low in the sulcal walls, matching predictions based on the orthogonal component of the electric field. The excitation in the sulcal cortical surface was consistent with the well-established columnar neuronal orientation and functional organization of the cortex and functional imaging studies^40,57^. Furthermore, the excitable area of the L5 PNs were wider in the postcentral gyrus compared to the precentral gyrus. This might be due to the severe curvature in the precentral gyrus or the thinner cortex on the postcentral side such that the neurons were smaller than those in precentral gyrus. This suggests an additional important factor of cortex geometry for TMS next to neuron placement and coil orientation^30^. The L3 PNs had different morphology with shorter axons than the L5 PNs such that they were located completely within the gray matter. Similar to the L5 PNs, the coil orientation had a significant impact on the responses of the L3 PNs, but the precise patterns of the threshold maps differed between the L3 and L5 PNs. As hypothesized by Day et al., here the proximity to the coil played an important role^58^, as L3 PNs in the gyral crown and the upper parts of the sulcal wall were predominantly activated.

The neural response to TMS is composed of a direct (D) and several indirect (I) waves. The D-wave is thought to be produced by direct activation of L5 PNs as we have modeled it here and is followed by I-waves that are thought to be generated by synaptic excitation and/or re-excitation of L5 pyramidal cells with longer latencies^58,59^, presumably via pyramidal cells in superficial cortical layers L2 and L3. According to Di Lazzaro et al. (2004), at the lowest stimulation intensity to evoke neuronal responses, an I-wave is elicited, and with increasing stimulation intensity, the earlier, small D-wave is produced^30^. This indicates that thresholds for eliciting I-waves are lower than those for eliciting D-waves^60^. In this work, we explored the excitation thresholds of both L3 and L5 PNs and found that the percentage of excited neurons for a stimulation intensity at the motor threshold was about 2 times higher for L3 PNs than for L5 PNs. Furthermore, the activation of the L3 PNs was consistently higher than that of the L5 PNs for the full range of stimulation intensities. The lower stimulation thresholds of the L3 PNs are consistent with lower stimulation intensities required to produce I-waves^49,61^, and the higher stimulation intensities required to produce D-waves.

The highest percentage of excited PNs was observed at +135 degrees and the excited regions were focused on the postcentral gyrus. The base coil orientation induced the lowest percentage of activated PNs, but as shown in Fig. 4 the precentral gyrus was targeted better than for other coil orientations. Thus, to activate the precentral gyrus, the base coil orientation is recommended by our model, congruent with previous research^62,63^, and +135 degrees should be ideal to stimulate the postcentral gyrus.

The question of the precise initiation site of action potentials is a central issue for understanding the physiological effects of TMS. According to our study, the dominant initiation site leading to action potentials is the axon initial segment in both L5 and L3 PNs. This is consistent with previous studies arguing that action potentials giving rise to the D-wave might be initiated close to the soma and/or axon initial segment^64,65^. In addition, L5 PN action potentials were also initiated at the axon where it crosses the boundary between gray matter and white matter, where tissue conductivity changes abruptly^48^. However, it will be important to verify these results with more realistic axon models.

There are several limitations in our modeling study. A first limitation is related to neuronal properties and morphologies. The L3 and L5 PNs were taken from cat visual cortex due to the lack of models for most human cortical cell types. However, despite the uncertainty with regard to properties of PNs, we produced results matching both experimental studies and other computational studies^23,25^ that incorporated the same models of PNs.

While we observed the stimulation of neural activity in the superficial cortex nearby the coil, TMS might also affect deep brain areas that cannot be stimulated directly. This can be explained on the basis of the propagation of action potentials along white matter fiber tracts. Recent studies modeled tractography-based white matter fiber tracts using diffusion tensor imaging (DTI) and observed activation of axon bundles^9,11,12^. Compared to fiber tracts in previous modeling, we modeled straightly stretched axons of L5 PNs inside the WM. Due to such limitations, the axons inside the WM occasionally passed through protruding parts of GM. Notwithstanding this intersection could affect the neuronal responses such as the action potential initiation or thresholds, most PNs initiated action potentials at the axon initial segment and coil orientation dependency observed in threshold maps was consistent with observations in previous studies. Further developments in tractography may improve detailed neuronal models and may lead to a deeper understanding of the TMS-induced brain activity propagations from the superficial cortex to distant brain regions.

Another limitation is that the reconstructed PNs were synaptically isolated. For the L5 cells this means that we basically studied the generation mechanism of D-waves. The activation of L3 cells could be seen as a proxy for the generation of I-waves. A logical next step is to synaptically couple L3 and L5 cells as done in a recent model for the generation of D and I-waves using L5 PNs that were contacted by a pool of excitatory and inhibitory layer 2 and 3 neurons^49^. This model successfully reproduced various characteristics of I-waves and highlighted the importance of the complex morphology of the L5 PNs for the generation of I-waves. An improvement would be to use the anatomical information on the activation of PNs as modeled here, as we found a clear difference in the threshold maps for L5 and L3 PNs based on their morphology. Therefore, in future work, we plan to incorporate synaptic connections between L3 and L5 PNs. We hope that this will bring us one step closer to a detailed understanding of the mechanisms through which TMS activates cortical circuits, paving the way for more precise and effective application of TMS based on individual brain morphology in clinical and basic research settings.

